# Phylogenomic Synteny Network Analysis Reveals an Ancient MADS-Box Transcription Factor Tandem Duplication and Lineage-Specific Transpositions

**DOI:** 10.1101/100990

**Authors:** Tao Zhao, Rens Holmer, Suzanne de Bruijn, Harrold A. van den Burg, M. Eric Schranz

## Abstract

Conserved genomic context (or synteny) provides critical information for comparative evolutionary analysis, such as the inference of ancient polyploidy events, recurrent genomic rearrangements across species and gene ancestry. With the increase of sequenced and assembled plant genomes, we now have the opportunity to use synteny to analyze the dynamics of gene family expansion and contraction across broad phylogenetic groups. Here we present an integrated approach to organize plant kingdom-wide gene synteny networks using *k*-clique percolation. As an example, we analyzed the gene synteny network of the MADS-box transcription factor family based on fifty-one completed plant genomes. We conclude from two massive gene clusters that one of the two Type II MADS-box gene clades evolved from an ancient tandem gene duplication likely predating the radiation of seed plants, which then expanded by polyploidy events and sub-functionalization. This gene clade now contains key regulators of major phenotypes of angiosperms including flower development. Moreover, we find lineage-specific gene clusters derived from transposition events. For example, lineage-specific clusters in the Brassicales containing genes that are well-known for their function in controlling flower morphology (*AP3* and *PI*). Our phylogenomic synteny network approach can be applied to any group of species to gain new insights into the evolution and dynamics of any set of genes.

## INTRODUCTION

Correlated gene arrangements (synteny) can be maintained across hundreds of millions of years and provides critical information about conserved genomic context and the evolution of genomes and genes. For example, the well-known “Hox gene cluster” that regulates the animal body plan is largely preserved across the animal kingdom (Lewis, 1978; Krumlauf, 1994; Ferrier and Holland 2001). Synteny data is widely used to establish the occurrence of ancient polyploidy events, identify chromosomal rearrangements, to examine gene family expansions and contractions and to establish gene orthology (Sampedro et al., 2005; Tang et al., 2008a; Jiao and Paterson, 2014). Conserved synteny likely reflects important relationships between genomic context of genes both in terms of function and regulation and, thus, is often used as a proxy for the conservation or constraint of function (Dewey, 2011; Lv et al., 2011). An investigation of the syntenic relationships across a wide range of species can be used to address fundamental questions on the evolution of important gene families, such as those fundamental to the regulation of developmental pathways. For example, the origin of morphological novelty has been linked to the duplication of key regulatory transcription factors such as the Hox-genes in animals and the MADS-box genes in plants (Alvarez-Buylla et al., 2000; Airoldi and Davies, 2012; Soshnikova et al., 2013).

To systematically investigate, sort, and visualize all possible microsyntenic (small regions of synteny) relationships between all gene family members across many species is challenging. This is due to the ubiquity of ancient and recent polyploidy events, as well as smaller scale events derived from tandem and transposition duplications (Lynch and Conery, 2000; Bowers et al., 2003; Tang et al., 2008a; Schranz et al., 2012). For example, the ParaHox and NK gene clusters important for animal body plan development are likely derived from an ancient duplication of the Hox gene cluster (Brooke et al. 1998; J Garcia-Fernàndez 2005). Also, even the well-known Hox-cluster is found be dispersed or “broken-up” in some animal lineages (Lemons and McGinnis, 2006; Duboule 2007; Albertin et al. 2015). Critical regulators of plant development, such as some MADS-box genes, are also derived from ancient duplication events (Ruelens et al., 2013).

In plants, the MADS-box genes are the major transcription factors regulating the specification of floral organs (e.g. the ABC(DE) model), reproductive organ identity and other traits (Theissen 2001; Becker and Theissen, 2003; Smaczniak et al., 2012). This multi-gene family has undergone extensive duplications leading to complicated relationships of orthology, paralogy, and functional homology (Jaramillo and Kramer, 2007). In the model plant *Arabidopsis thaliana*, there are 107 MADS-box genes that are derived from multiple gene duplication events (Martinez-Castilla and Alvarez-Buylla, 2003; Parenicova et al., 2003). The MADS-box genes were divided into two major clades, termed Type I and Type II. The Type II lineage is further divided into the MIKC^C^- and MIKC*-types (Henschel et al., 2002). The function and evolution of MADS-box genes have been extensively studied, especially the MIKC^C^-types (for review see Smaczniak et al., 2012). Synteny data of MADS-box genes has partially been utilized to investigate the ancestral genetic composition of the B- and C-function (Causier et al., 2010) and the A- and E-function genes (Ruelens et al., 2013; Sun et al., 2014). However, these studies included limited set of species (<10) in their synteny comparisons, the results were displayed as linear pair-wise comparisons (as exampled in Figure 1a), and more importantly, complete syntenic data for the entire MADS-box gene family yet not feasible in one study. When more genomes for a gene family are simultaneously analyzed at both the microsynteny and gene similarity level, it becomes increasingly more difficult to organize and display the syntenic relationships that emerge.

**Figure 1.**
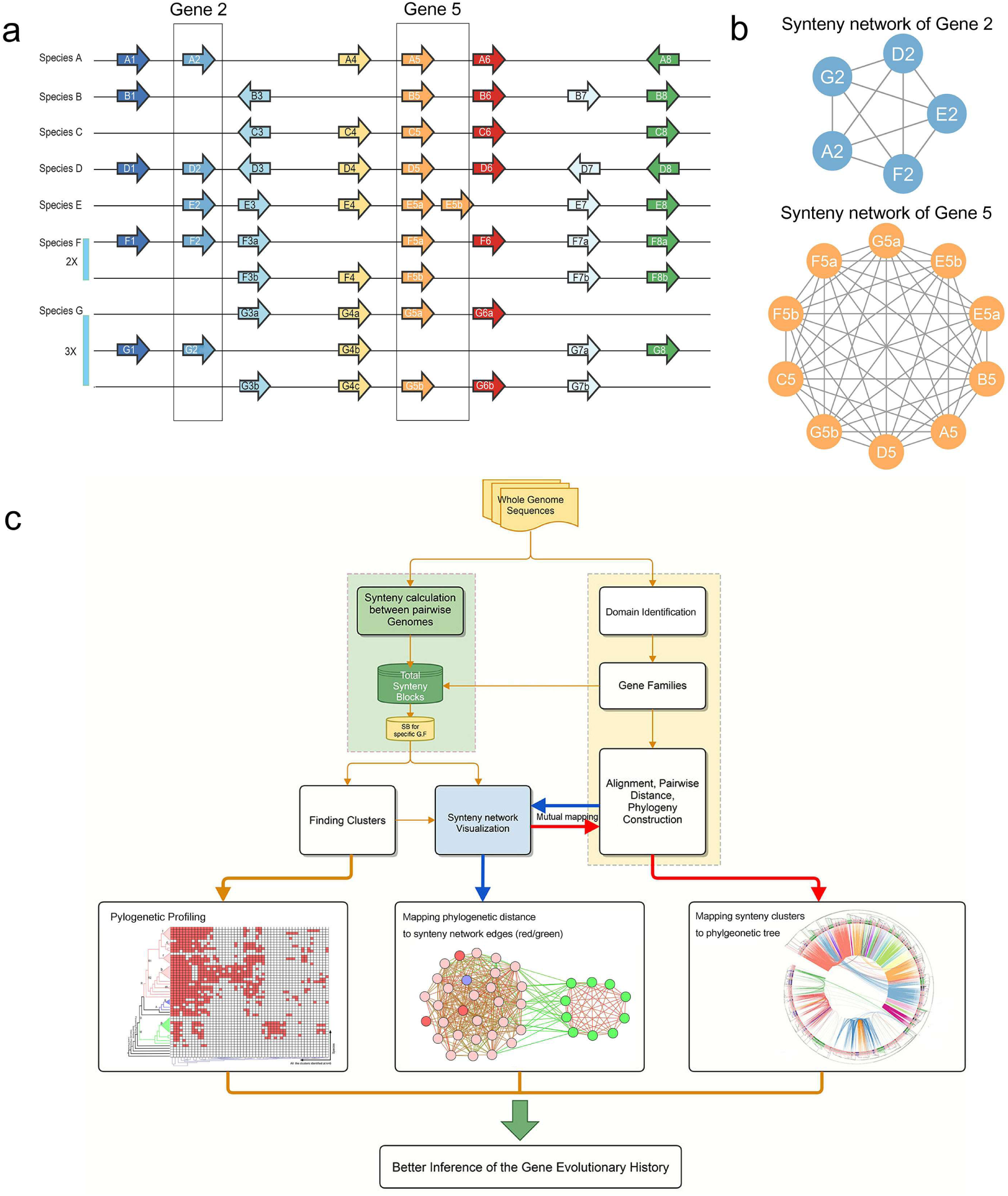
Principle and workflow of the synteny network analysis. **(a)** Simulation of a typical pair-wise based syntenic alignment sketch across seven species (A-G) for which species F has undergone a Whole Genome Duplication (WGD = 2x) and species G a Whole Genome Triplication (3x). Gene 2 and gene 5 are boxed respectively as examples for a further network view of synteny relationships. **(b)** Undirected, unweighted, circular network view organizing pairwise syntenic connections, or edges, of target genes (in this case, gene 2 and gene 5) across the seven species from (a), nodes represent syntenic genes and edges (lines) represent a syntenic connection between two nodes. **(c)** The workflow pipeline for synteny network. Annotated whole genome sequences enter the pipeline and are used in two parallel modules. The first, light green rectangle, represents the analysis of pairwise synteny calculation (synteny block detection) that generates a syntenic network database (dark green). The second, light yellow rectangle, represents a phylogenetic analysis including gene family identification, sequence alignment and gene tree construction. Synteny blocks for a specific gene family (yellow) can be obtained using the output from gene family identification. The network data is then processed by clustering and visualization, sub-networks from clustering are used for phylogenetic profiling (bottom left) to depict node composition at species level. Examples of mutual mapping between syntenic connections and phylogeny: Phylogenetic pairwise distance can be indicated on syntenic network, using edge color from red to green (bottom middle); syntenic relationships and communities can be inferred over the phylogenetic tree, using inner colored links (bottom right).

Here, we present a pipeline for gene ancestry based on synteny and that applies network analysis methodology to investigate broad microsynteny patterns. By displaying shared synteny between the multiple genomes as a network of connected genes (Figures 1b and 1c), we can tap into the wealth of existing network analysis and visualization tools (Lancichinetti and Fortunato, 2009; Fortunato, 2010). For example, for network clustering, the *k*-clique percolation and other methods have to date provided some of the most successful ways to consider community overlap (Palla et al., 2005; Porter et al., 2009). Our network approach facilitates detection and visualization of both intra- and interspecies syntenic connections across many species and multiple ancient polyploidy events. This integrated analysis revealed previously undetected evolutionary patterns of gene duplication and shared gene ancestry for the MADS-box transcription factor gene family during angiosperm evolution.

## RESULTS

### Overview of the synteny network pipeline

To explain our approach, we use the plant MADS-box gene family as an example. The analyzed genomes included green algae, mosses, gymnosperms, and angiosperms; in total fifty-one plant species (Supplemental Table 1). We built a database that contained links of syntenic gene pairs within syntenic genomic blocks based on the tool MCScanX (Tang et al., 2008b; Wang et al., 2012). This database, with all-pairwise syntenic blocks from these 51 species, contained 921,074 nodes (i.e. genes that were connected by synteny with another gene) and 8,045,487 edges (i.e. pairwise syntenic connections). To extract the network of the MADS-box gene family from this entire gene synteny network, we then identified the MADS-box genes using HMMER3.0 screening the predicted coding sequences from all test genomes. This yielded a total of 4,221 MADS-box genes (Supplemental Table 2, sheet 1). This allowed us to extract the synteny network of the MADS-box gene family from the whole synteny network. We then visualized this MADS-box gene synteny subnetwork using Cytoscape and we applied a community detection algorithm (application of clustering methods), and phylogenetic profiling (Figure 1c). In this way, we obtained syntenic networks with a total of 3,458 MADS box gene nodes, which are linked by 25,500 syntenic edges (Figure 2b; Supplemental Table 2, sheet 2).

**Figure 2.**
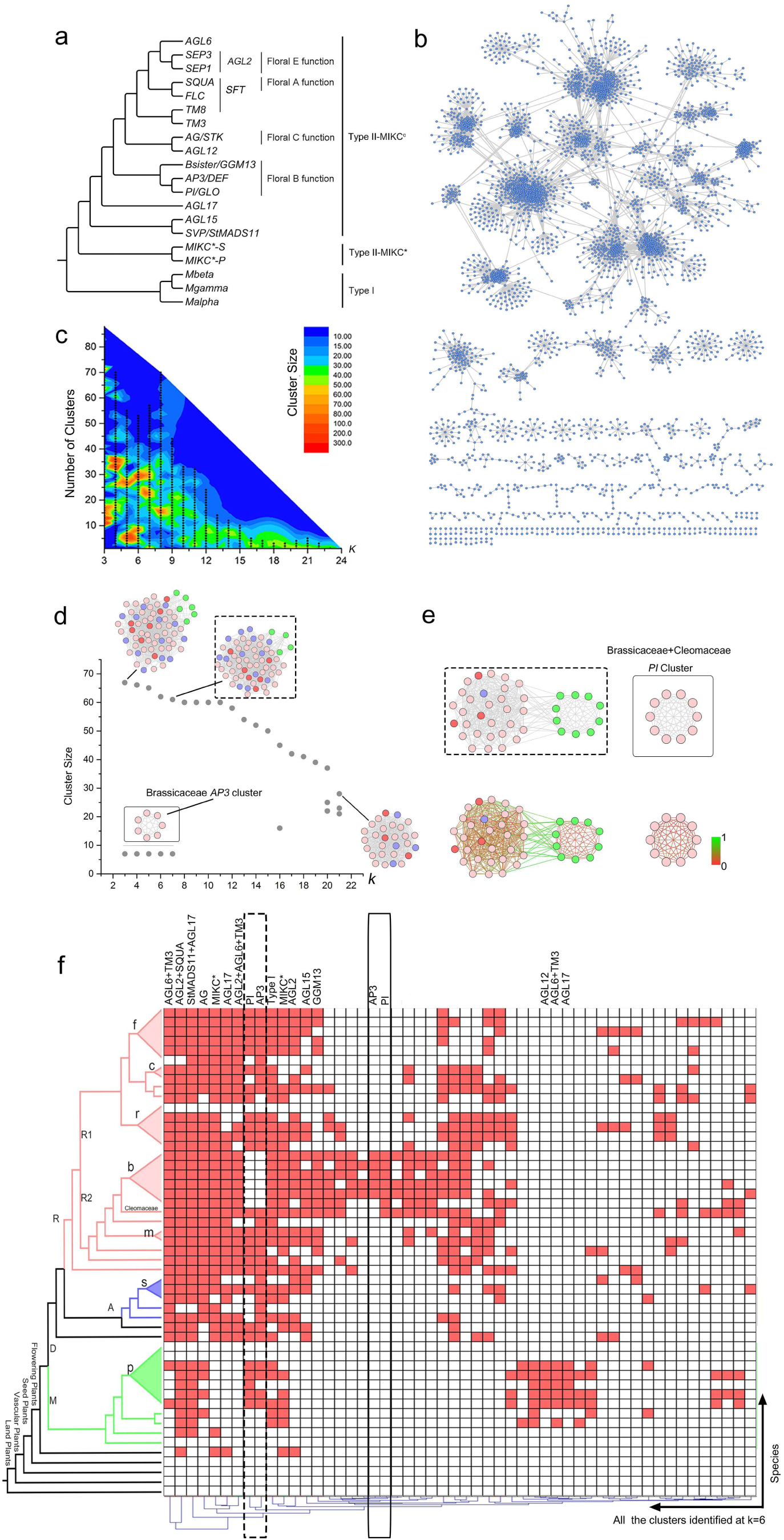
Synteny network analysis outputs of the plant MADS-box gene family from 51 target genomes. **(a)** The consensus phylogenetic tree showing relationships of the subfamilies of the MADS-box gene family based on a number of previous studies. Note that for convenience in this study, clades containing *SQUA*-, *FLC*-, and *TM8*-like genes are named *SFT*. **(b)** A network view depicting all the syntenic relationships for the entire MADS-box gene family from fifty-one species (undirected, unweighted graph). The nodes represent individual MADS-box genes, edges represent syntenic connections between the nodes derived from the syntenic blocks. **(c)** Number of clusters (*k*-cliques) and cluster (clique sizes) under a certain *k* from (b). **(d)** An example of the size change of the *AP3* subfamily syntenic *k*-cliques across k values. Node colors represent rosids (light pink), asterids (blue), monocots (green), and nodes belonging to *Amborella trichopoda*, *Vitis vinifera*, *Beta vulgaris*, and *Nelumbo nucifera* are in red. **(e)** *PI* clusters, where the edge colors represent the phylogenetic distance (bottom right). The green edges are between monocot and eudicot species. **(f)** Phylogenetic profile of all the *k*-cliques at *k* = 6. Letters on the phylogenetic tree represent: f (Fabaceae), c (Cucurbitaceae), r (Rosaceae), b (Brassicaceae), m (Malvaceae), s (Solanaceae), p (Poaceae), D (Eudicots), M (Monocots), A (Asterids), R (Rosids), R1 (Eurosids I), and R2 (Eurosids II). Red colored field indicates presence of at least one syntelog of the MADS-box gene in that species. The phylogenetic profiling approach led to the identification of highly conserved communities compositions such as “*AGL6+TM3*” and “*AGL2+SQUA*”; transposed synteny networks of *PI* and *AP3* across Brassicaceae; also monocots specific ones like “*AGL12*”, “*AGL17*”, and “*AGL6+TM3*”. Corresponding abbreviations of the species names are shown on the right side. The dotted box and the line box represent the corresponding networks in (d) and (e).

For clarity, we have summarized the current notion of evolutionary relationships between the major clades of the MADS-box gene family (Figure 2a) based on numerous studies (Becker and Theissen, 2003; Martinez-Castilla and Alvarez-Buylla, 2003; Nam et al., 2003; Nam et al., 2004; Nam et al., 2005; Smaczniak et al., 2012; Gramzow and Theissen, 2013; Kim et al., 2013; Ruelens et al., 2013; Gramzow et al., 2014; Sun et al., 2014; Yu et al., 2016). Note that we have named the hypothesized common ancestral genes of ***S****QUA*-, ***F****LC*- and ***T****M8*- like genes as *SFT* for convenience (the abbreviation *SFT* will be used later in this study).

One important characteristic of synteny-based analyses is the fact that closely related species have increased numbers of syntenic regions that span large genomic regions (e.g. species within the Brassicaceae family), which provides highly connected nodes in the network, while more remote species (e.g. comparisons to the basal angiosperm *Amborella trichopoda*) show a lower number of syntenic regions with less evidence of synteny but they hold crucial evolutionary evidence. We here use CFinder to identify structure in the MADS-box synteny network using the *k*-clique percolation method (Palla et al., 2005; Palla et al., 2007). *K*-clique corresponds to a fully connected sub-selection of *k* nodes (e.g. a *k*-clique of *k* = 3 is equivalent to a triangle). Two *k*-cliques are considered adjacent and thus form a *k*-clique-community if they share *k*-1 nodes (Derenyi et al., 2005; Palla et al., 2005). In this study, all cliques of size *k* = 3 to *k* = 24 were detected for the MADS-box gene synteny network using CFinder, as were the number of *k*-clique-communities under each *k*-clique (Figure 2c). Each of the community sizes under a certain *k* are shown (Figure 2c), which quantifies the strength of the syntenic connections within and between subfamilies across species. For example, the *AP3*-like genes of monocots species (green nodes) are only part of a community at relatively low *k* values (*k* < 8) (Figure 2d). Hence, the *k* values enable us to optimize cluster sizes in a way that minimizes weakly adhered nodes while retaining as much information as possible.

We then calculated the protein sequence pairwise distances (Supplemental Figure 1a) and visualized these distances using an edge color key (from red to green; Figures 2e, 3c and 3d). This color key highlights the degree of sequence divergence between syntenically connected genes. For example, two separated *Pistallata* (*PI*) communities were found from the previous *k*-clique percolation step (Figure 2e top panel); one is shared by most monocot and eudicot species (*k* = 6, community size 40, Figure 2e left panel), while the other one is shared by species in Brassicaceae and Cleomaceae (*k* = 6, community size 10, Figure 2e right panel). The sequence divergence of the nodes can be inferred upon mapping the edge color key (Figure 2e bottom panel). For such genes linked by green edges, it can occur that they group in a single synteny community, but spread over distinct remote clades in a phylogenetic analysis.

**Figure 3.**
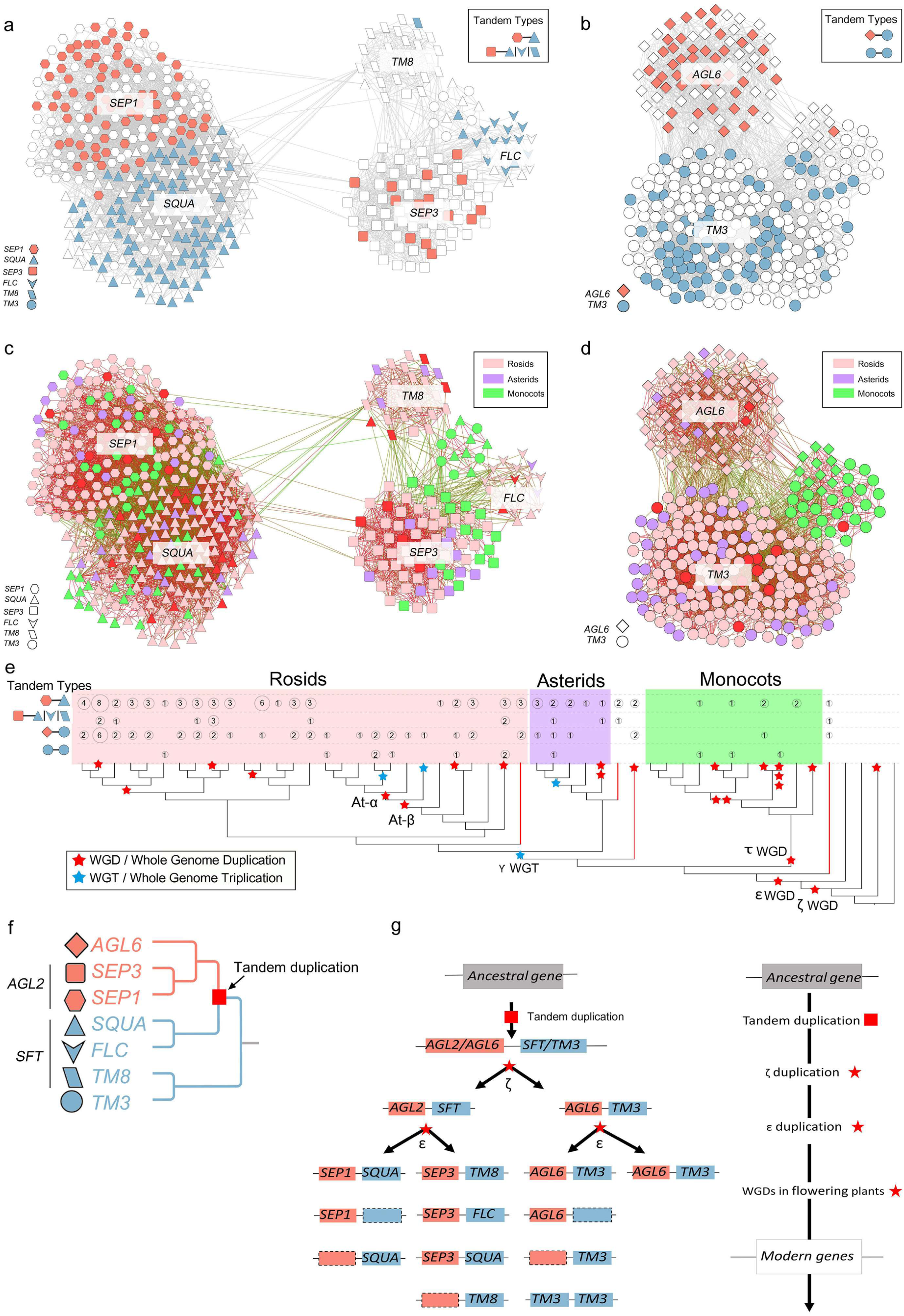
Evolutionary history of the major MIKC^c^-type MADS-box gene super-clade. **(a)** Synteny network of *AGL2* (*i.e. SEP1, SEP3*)-, *SQUA*-, *TM8*-, and *FLC*-like gene clusters. Tandem gene arrangements of *SEP1-SQUA*, and *SEP3-SFT* are highlighted; *SFT*= *SQUA/FLC/TM8*. **(b)** Synteny network of the *AGL6*- and *TM3*-like gene clusters. Tandem gene arrangement of *AGL6-TM3* and *TM3-TM3* are highlighted. **(c)** Synteny network of *AGL2*-, *SQUA*-, *TM8*-, and *FLC*-like gene clusters, node color represents species as defined in Figure 2d, edge color represent the phylogenetic distance as defined in the main text. Nodes/genes from different clades are indicated by node shapes. **(d)** Synteny network of the *AGL6-* and *TM3*- like gene clusters. Nodes/genes from different clades are indicated by node shapes. **(e)** Summary of the occurrence of the tandem gene arrangements from (a) and (b) across the species analyzed. Top two rows quantify the number of *SEP1-SQUA* and *SEP3-SFT* tandems in (a) respectively, while the third and fourth rows sum up the number of *AGL6-TM3* and *TM3-TM3* tandems in (b). Polyploidy events are highlighted on the species tree, where blue indicates genome triplication and red indicates genome duplication, a same tree with full species names and known duplication events can be found in Supplemental Table 1. **(f)** Mapping tandem arrangements inferred from the summary of synteny networks in (e) to the consensus phylogeny. **(g)** Proposed evolutionary scenario for the origin of the MADS-box gene family clade in (f).

A clique size *k* = 3 to 6 has been found to best approximate the true number of communities (Derenyi et al., 2005; Palla et al., 2005; Porter et al., 2009; Xie et al., 2013). In this study, fifty-three communities (distinct clusters) were obtained for *k* = 6 (Figure 2c and Supplemental Table 3) and thus were used for phylogenetic profiling (Figure 2f). Each column depicts a syntenic occurrence for a certain MADS-box gene in a particular plant species. Thereby the presence/absence of syntenic gene clusters across the 51 analyzed taxa are represented by their respective phylogenetic profiles to determine and infer evolutionary patterns (e.g. conserved vs. family-specific transpositions (such as the boxed *PI* and *AP3* clusters)). It is worth mentioning that for two monocot species, *Triticum urartu* (wheat) and *Hordeum vulgare* (barley), we did not find any syntenic regions for any of their MADS-box genes with other plant genomes at any *k* (Figure 2f), which is likely due to the fragmented genome assemblies of these two grass species.

As a highlight of the above pipeline and analysis approach, we are able to present new insights into the MADS-box gene family based on synteny network cluster analysis, which we describe in greater detail below.

### Angiosperm-wide conserved *SEP1-SQUA* and *SEP3-SFT* tandems

Based on our global analysis of the MADS-box genes with various *k* values, we were able to draw inferences about the evolution of particular gene clades. The largest community (475 nodes) was found when *k* = 3 was used (Figures 3a and 3c). The entire network can be divided into two parts: on the left are *SEP1*-, and *SQUA*-like genes, while on the right are the *SEP3*-, *SQUA*-, *FLC*-, and *TM8*-like genes (Figures 3a and 3c). The *SEP1*-, and *SQUA*-like genes are highly interconnected between and within genomes (Figures 3a and 3c) with syntenic orthologs being present for both genes in a wide-range of angiosperm species including the basal angiosperm *A. trichopoda*, monocots and eudicots. Further investigation revealed that the *SEP1*- and *SQUA*-like genes are arranged as tandem gene duplicates in most angiosperm species (Figures 3a and 3e, Supplemental Table 4), suggesting that this duplication occurred prior or at the origin of the angiosperms. For example, there is one *SEP1-SQUA* tandem gene arrangement in *A. trichopoda* and three such tandem gene arrangements are identified in grapevine following the gamma triplication in eudicots (Figures 3a and 3e, Supplemental Table 4).

This ancient *SEP1-SQUA* tandem gene arrangement, as revealed by the angiosperm-wide synteny network analysis, is in agreement with other studies where the *SEP1-SQUA* tandem gene arrangement was found in eudicots (Ruelens et al., 2013). Sun and co-workers also noted that most *AP1*-like genes (a subclade of the *SQUA*-like genes) and *SEP1*-like genes were tightly linked as genomic neighbors since the split of the basal eudicots (Sun et al., 2014).

In addition, from the right half of the network in Figures 3a and 3c, our data show that most eudicot *SEP3*-like genes group as a distinct cluster, which is relatively loosely connected (green edges) to nodes representing monocot *SEP3*-, *SQUA*-, and *TM3*-like genes (monocot nodes are in green). This *SEP3*-like gene cluster is also connected to the eudicot *FLC*- and *TM8*-like genes (Figures 3a and 3c). Based on synteny analysis, it was previously suggested that these *SEP3*- and *FLC*-like genes originate from an ancient tandem gene duplication (Ruelens et al., 2013). We also identified this *SEP3-FLC* tandem gene arrangement in eight eudicots species (Figures 3a and 3e, Supplemental Table 4). However, the *SEP3-FLC* tandem gene arrangement is found less often than the *SEP1-SQUA* tandem gene arrangement. Besides this, we found that in *A. trichopoda* the *SEP3* and *TM8* homologs are also arranged in tandem (*SEP3-TM8*). In two monocot species, *Oryza sativa* and *Sorghum bicolor*, *SEP3* and *SQUA* are found in tandem (Figures 3a and 3e, Supplemental Table 4). *TM8* was first identified from *Solanum lycopersicum* (Pnueli et al., 1991), and this clade of genes has been reported to have undergone independent gene loss in different lineages based on phylogenic analyses (Becker and Theissen, 2003; Gramzow and Theissen, 2013). According to the consensus phylogeny based on studies by others (Figure 2a), the TM8-like genes are close to *TM3*-like genes and they both appear to share a common origin with the *AGL6*-, *AGL2*-, *SQUA*-, and *FLC*-like genes (Figure 2a).

Our synteny analysis reveals a more broadly conserved, and thus potentially more ancient, tandem gene duplication that involves the last common ancestor of all *SEP3*- and *TM8*-like genes. Considering that *TM8*-like genes were already present in the last common ancestor of extant seed plants (Gramzow et al., 2014), it is likely that the *SEP3-TM8* tandem is more ancestral than the *SEP3-FLC* tandem. Hence, the *FLC*-like genes could be derived from a *TM8* homolog in an ancestral plant. According to the network structure and gene copy number of the *SEP3*-, *FLC*- and *TM8*-like gene clusters, we hypothesize that after the split with *A. trichopoda* the *SEP3*- and *TM8*-like genes likely do not appear a tandem gene pair within one species and after WGDs *TM8*-like homologous tend to be lost from the tandem. This means that the *SEP3-TM8/FLC* tandem gene pair is more variable than the *SEP1-SQUA* tandem gene pair. Previous work suggested that these two tandems derive from a whole-genome or large scale segmental duplication in a common ancestor of the angiosperms (Ruelens et al., 2013). Indeed, we found both the *SEP1-SQUA* and *SEP3-TM8* tandem gene pair in the basal angiosperm *A. trichopoda* (Figure 3e, Supplemental Table 4). Hence, the duplication of these two tandems via a WGD very likely occurred around the ε WGD event, derived from one ancestral tandem gene pair of *AGL2-SFT* (*AGL2* = *SEP*, *SFT* = *SQUA/FLC/TM8*, Figure 3g) in a common ancestor of the angiosperms (Jiao et al., 2011).

Finally, the *FLC* homologs that are found in the genomes of Brassicaceae and Cleomaceae species do not group with the angiosperm-wide *FLC* cluster, instead they form their own independent cluster (Supplemental Figure 1b). This suggests that there was a gene transposition of the ancestral *FLC* gene in the Brassicales lineage after the split of the early branching Papaya, potentially near the time of the At-β WGD (Edger et al., 2015).

### Angiosperm-wide conserved *AGL6-TM3* tandems

According to the phylogenetic analyses of the MADS-box genes, the *AGL6*-like genes form a sister clade to the *AGL2*(*SEP*)-like genes (Figure 2a) in both angiosperm and gymnosperm (Becker and Theissen, 2003; Zahn et al., 2005; Ruelens et al., 2013; Yockteng et al., 2013; Sun et al., 2014).

In this study, we observed another large synteny community (*k* = 3, community size: 305) that contains the *AGL6*- and *TM3* (*SOC1*)-like genes (Figures 3b and 3d). Like the *SEP1-SQUA* and *SEP3-SFT* tandems in Figure 3a, we found prevalent presence of *AGL6-TM3* tandems and also some *TM3-TM3* tandems ( Figures 3b and 3e, Supplemental Table 4). For example, in basal angiosperms there is one *AGL6-TM3* tandem gene pair in *A. trichopoda* and two such tandems in *Nelumbo nucifera* likely due to the most recent WGD this species experienced (Ming et al., 2013; Wang et al., 2013) (Figure 3e, Supplemental Table 4). In grape (*V. vinifera*) we also found two *AGL6-TM3* tandems (Figure 3e, Supplemental Table 4), these could originate from the *γ* whole genome triplication (WGT), after which one tandem lost its *AGL6* locus. This is in agreement with a previous study (Vekemans et al., 2012). Like *V. vinifera*, *T. cacao* has not undergone any additional WGD after the *γ* WGT and also in this genome two *AGL6-TM3* tandems remain.

Besides the prevalent *AGL6-TM3* tandem gene arrangement, we also found *TM3-TM3* tandems in ten species (seven eudicots species and three monocot species) (Figure 3e, Supplemental Table 4). Hence, the network in Figures 3b and 3d has overall more *TM3* genes than *AGL6* genes.

### Evolutionary history of the major MIKC^c^-type MADS-box gene super-clade

The general notion is that the *AGL6*- and *AGL2* (*SEP*)-like genes are close homologs (Figure 2a) and it has been hypothesized that the combined ancestral gene of the *AGL6-* and *AGL2*-like genes was duplicated in a common ancestor of the seed plants (Spermatophytes) (Zahn et al., 2005; Kim et al., 2013), probably as a result of the ζ WGD (Jiao et al., 2011). By interpreting the synteny networks, we found strong evidence of *SEP1-SQUA*, *SEP3-SFT*, and *AGL6-TM3* tandems (Figures 3a, 3b, and 3e), and also evidence of monocot *TM3*-like genes connected to *SEP3*-, *SQUA*-, and *TM8*-like genes (Figure 3c). This enabled us to deduce the deep genealogy and to propose an evolutionary diagram that depicts how one ancestral locus that predates the last common ancestor of all seed plants has given rise to a large MADS-box gene clade with many subfamilies in angiosperms, which includes the *AGL2*-, *AGL6*-, *SQUA*-, *TM3*-, *TM8*-, and *FLC*-like gene clades (Figures 3e-3g). It can be inferred that in the last common ancestor of seed plants a gene tandem was already present that corresponds with the current *AGL2/AGL6-SFT/TM3* tandem gene arrangement (Figures 3f and 3g). The ζ WGD (shown in Figure 3e) that occurred shortly before the radiation of the extant seed plants (Jiao et al., 2011) is likely causal to the duplication of this original tandem gene pair, after which the *AGL2*- and *AGL6*-like genes then diverged, as well as the *SFT*- and *TM3*-like genes (Figure 3g). A subsequent more recent WGD (the ε event), which occurred prior to the diversification of the extant angiosperms (Jiao et al., 2011), allowed then the emergence of the *SEP1*- and *SEP3*-like genes from the ancestral *AGL2* locus, as well as the *SQUA*-, *TM8*-, and *FLC*-like genes from the ancestral *SFT* gene. During that same period only one copy of the *AGL6-TM3* tandem was retained from the ε WGD (Figure 3g).

However, we did not find supporting tandems from the gymnosperm species *Picea abies* in the synteny network data to further prove our hypothesis. This is probably due to the high proportion of transposable elements in this gymnosperm genome that strongly reduced its synteny with other plant genomes (Nystedt et al., 2013).

### Distinctive *AP3* and *PI* clusters in Brassicaceae and Cleomaceae

Another interesting finding from our synteny approach in the MADS-box family involves the *AP3* and *PI* genes, which belong to MADS-box floral B-class genes. These two genes are important for petal and stamen specification (Jack et al., 1992; Goto and Meyerowitz, 1994; Jack et al., 1994; Zhang et al., 2013). We found that most *AP3* genes reside in a single cluster that comprises homologs of both eudicot and monocot species, including the basal angiosperm *A. trichopoda* and basal eudicot *N. nucifera*. However, the cluster lacks *AP3* homologs from Brassicaceae species (Figure 2d). Instead, the *AP3* genes from Brassicaceae form a separate cluster (except for *Aethionema arabicum*, *Aethionema AP3* gene was annotated on a scaffold with no other genes (gene name: *AA1026G00001*, highlighted in Supplemental Table 2, sheet 1)). A very similar picture emerges when analyzing the *PI* genes: *PI* homologs from six Brassicaceae species group together with one *PI* gene from *Tarenaya hassleriana* (a Cleomaceae species), while the *PI* homologs from most other species form a second distinct cluster (Figures 2e and 2f). These patterns suggest that in the case of both *PI* and *AP3*, a transposition or genomic rearrangement event resulted in a unique genomic context for these genes in Brassicaceae. Since one Cleomaceae *PI* gene belongs to the Brassicaceae *PI* cluster (Figure 2e right panel) but the Brassicaceae *AP3* cluster does not contain any Cleomaceae *AP3* gene (Figure 2d solid line boxed cluster and Figure 2f), we speculate that the different syntenic position of Brassicaceae *AP3* and *PI* genes are initially the result of the genomic rearrangement from the At-α and At-β WGDs, respectively (At-α and At-β WGDs are shown in Figure 3e).

In the model plant *A. thaliana*, the AP3 and PI proteins form obligate heterodimers and bind CArG-box cis-regulatory sequences in promoter elements (Riechmann et al., 1996; Yang et al., 2003). By such heterodimerization and/or homodimerization, a wide array of potential protein-protein interactions can evolve that can lead to the highly diverse flower morphologies in angiosperms (Lee and Irish, 2011; Melzer et al., 2014; Bartlett et al., 2016). Interestingly, it is well known that the Brassicaceae species have rather uniform, or canalized, flowers (typical cross arrangement of the four petals). However, in its closest sister family Cleomaceae, which diverged from each other only ~ 38 million years ago (MYA) (Schranz and Mitchell-Olds, 2006; Couvreur et al., 2010), more diverse floral morphologies are observed (Patchell et al., 2011). In this study, we found unique synteny pattern of *T. hassleriana* B-genes that is consistent with earlier findings (Cheng et al. 2013). One *T. hassleriana PI* gene sits in the cluster shared with most other eudicots and monocot species, while the other *T. hassleriana PI* gene sits in the cluster shared with Brassicaceae species (Figures 2e and 2f). This suggests that the At-β WGD gave birth to new genomic position for one *PI* gene, while another *PI* gene still kept the original synteny to other eudicot and monocot species in the last common ancestor of Brassicaceae and Cleomaceae. After that, most likely due to the At-α WGD, Brassicaceae species lost the *PI* gene syntenic to other eudicots and monocots species, in addition, led to a unique synteny for the Brassicaceae *AP3* genes (Figure 2d). Therefore, such distinctive syntenic networks of *PI* and *AP3* caused by genomic rearrangements as we found in Brassicaceae indicate that genomic context change may eventually have facilitated modified protein-protein interactions (through a change of promoter sequence and/or coding sequence), and potentially the canalization of the crucifer floral form.

### Synteny of *StMADS11-* and *AGL17-like* genes

Homologs of both the *StMADS11*- and *AGL17*-like gene families were recently shown to be essential in angiosperms (Gramzow and Theissen, 2015). In *A. thaliana*, the *StMADS11* gene clade is composed of two genes called *SVP* (*AGL22*) and *AGL24*. These two genes regulate the transition to flowering in *A. thaliana* (Hartmann et al., 2000; Michaels et al., 2003). In *A. thaliana*, there are four homologous genes that group in the *AGL17*-like subclade. Two of them, *ANR1* and *AGL21* were found to have key roles in the lateral root development (Zhang and Forde, 1998; Yu et al., 2014). The other two genes, *AGL16* and *AGL17*, play a role in the control of flowering (Han et al., 2008). Interestingly, we found that the synteny community that comprise the *StMADS11*- and the *AGL17*-like genes are moderately interconnected (Supplemental Figure 1c). This indicates that these two gene clades (*AGL17* and *StMADS11*), which both play key functions in plant development, potentially derive from a single ancestral locus that potentially duplicated due to a WGD event.

Specifically, in the *AGL17*-like genes cluster, genes from the basal angiosperm species *A. trichopoda*, the basal eudicot species *N. nucifera*, and most eudicot species are included. However, most monocot species *AGL17*-like genes were excluded (only one monocot node in *Musa acuminata*) (Supplemental Figure 1c). Instead the monocot *AGL17*-like genes form a specific cluster (Supplemental Figure 1d). This may be due to the ancient τ WGD shared by all monocot species (Jiao et al., 2014) (τ WGD was shown in Figure 3e). A previous study on the phylogeny of the *AGL17*-like genes also suggested a relative complex evolutionary history due to ancient gene duplications before the monocot-eudicot split (Becker and Theissen, 2003). Our data provide further evidence at the synteny level.

The *StMADS11*-like synteny network cluster (Supplemental Figure 1c top cluster) demonstrates that *SVP* (*AGL22*)- and *AGL24*-like genes are highly interconnected across most angiosperm species. This implies that also these genes are duplicates from a WGD. Importantly, the *A. thaliana SVP* gene was previously characterized as a floral repressor (Hartmann et al., 2000) and as a key gene for the regulation of the primary metabolism under drought conditions (Bechtold et al., 2016). In contrast, *AGL24* promotes flowering and can regulate the *SOC1* gene expression (Michaels et al., 2003; Torti and Fornara, 2012). Clearly, the *SVP* (*AGL22*) and *AGL24* clades have acquired novel potentially opposing functions since their emergence from a WGD duplication.

### Conserved Synteny found for MIKC*-type genes

The MIKC*-type genes form a monophyletic clade within the MADS-box genes (Alvarez-Buylla et al., 2000), with several of them being reported to play a major role in pollen development (Verelst et al., 2007a; Verelst et al., 2007b; Adamczyk and Fernandez, 2009). By phylogenetic analysis, expression and interaction data, this clade of genes was shown to have a conserved function throughout the evolution of vascular plant gametophytes (Kwantes et al., 2012; Liu et al., 2013). In *A. thaliana*, *AGL30*, *AGL65*, and *AGL94* form one monophyletic clade (MIKC*-P clade) and *AGL66*, *AGL67*, and *AGL104* form another clade (MIKC*-S clade) (Kofuji et al., 2003; Nam et al., 2004).

Using our synteny network, we found two highly connected networks that contain the angiosperm *AGL30*-, *AGL65*-, and *AGL94*-like genes (MIKC*-P clade) (Supplemental Figure 1e) and the *AGL66*-, *AGL67*-, and *AGL104*-like genes (MIKC*-S clade) (Supplemental Figure 1f), respectively (Supplemental Figure 1e). Both clusters encompass eudicots and monocot species, as well as the basal angiosperm *A. trichopoda.* However, the MIKC*-S cluster appears to have expanded in monocots, while homologs of *Nelumbo nucifera* are absent in this cluster (Supplemental Figure 1f). This means that both two MIKC* clades are broadly conserved across angiosperms. Interestingly, MIKC* protein complexes play an essential role in late pollen development in *A. thaliana* and the formation of this protein complex requires MIKC* proteins from both clades. For example, the AGL30 and/or AGL65 proteins from the P clade form heterodimers with AGL104 or AGL66, which both group with the S clade (Verelst et al., 2007a; Verelst et al., 2007b). This indicates that these two clades (gene clusters) have been functionally retained during angiosperm evolution.

### Scattered syntenic links for Type I MADS-box genes

The Type I MADS-box genes have very different evolutionary dynamics than the better-studied Type II MADS-box genes. For example, the Type I genes show a higher rate of gene birth-and-death (Nam et al., 2004) and gene duplication-transposition (Freeling et al., 2008; Wang et al., 2016). Also, the functions of Type I genes are less known. However, several Type I genes have been reported to play a role in female gametogenesis and seed development (Portereiko et al., 2006; Bemer et al., 2010). With our approach, we found two distinct clusters that contain Type I MADS-box genes (Supplemental Figures 1g and 1h). For example, the *PHERES1* (*PHE1*, *AGL37*) genes, which are regulated by genomic imprinting (Kohler et al., 2003), are in the same synteny network as *PHERES2* (*PHE2*, *AGL38*), *AGL35*- and *AGL36*-like genes, which all belong to the Mγ clade of the Type I MADS-box genes (Supplemental Figure 1g). Likewise, we found one cluster that contains genes from the Mα clade (Supplemental Figure 1h). Also, from both clusters, we can observe an overall higher sequence divergence rate as the degree of retained synteny between different genomes is reduced in both clusters independent of the distance between the plant species in these clusters, as evidenced by the green edges (Supplemental Figures 1g and 1h).

All pairwise syntenic relationships identified were further represented on a global phylogenetic tree of all MADS-box genes (syntenic and non-syntenic) (Figure 4a). The color of the connecting line indicates network communities at *k* = 3 (Figure 4b). The global comparison of synteny and phylogenetics (Figure 4) highlights the major differences between Type II clusters with a higher density of syntenic links and of Type I genes where syntenic links are sparser and jumpy (Figure 4a), indicating their unique evolutionary pattern, probably due to a higher frequency of gene duplication and transposition and weaker purifying selection (Nam et al., 2004). Figure 4 also conveniently conveys the ancient tandem duplications discussed above (e.g. *SEP1*-*SQUA*, *SEP3-SFT*, and *AGL6-TM3*).

**Figure 4.**
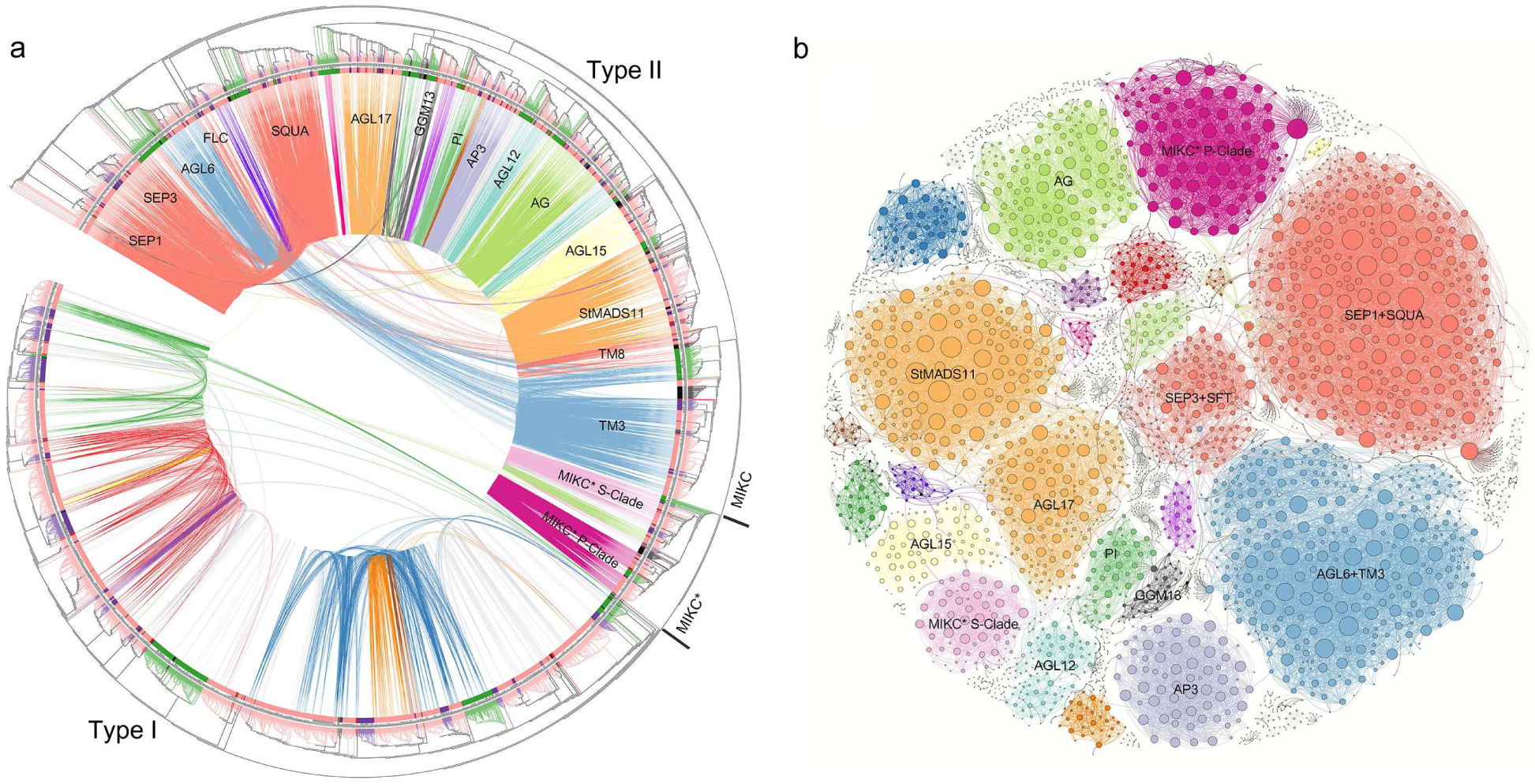
Phylogenic gene tree and synteny network of the plant kingdom MADS-box genes. **(a)** Maximum-likelihood gene tree for the MADS-box gene family and syntenic relationships among the genes. Boundaries of Type I, Type II, and MIKC- and MIKC*-Type II MADS-box genes are indicated on the tree. Terminal branches colors represent genes belonging to rosids (light pink), asterids (purple), monocots (green), genes belonging to *Chlamydomonas reinhardtii*, *Physcomitrella patens*, *Selaginella moellendorffii* and *Picea abies* are in black, and genes belonging to *Amborella trichopoda*, *Vitis vinifera*, *Beta vulgaris*, and *Nelumbo nucifera* are in red. Each connecting line located inside the inverted circular gene tree indicates a syntenic relationship between two MADS-box genes. The connecting lines are colored according to the discovered communities from (b). **(b)** Synteny network of the MADS-box gene family; communities were rendered based on the clique percolation method at *k* = 3. The size of each node corresponds to the number of edges it has (node degree).

## DISCUSSION

Syntenic analysis is an increasingly informative and robust approach to understand the evolutionary origin of genes and gene families. A typical analysis of synteny between two genomes (or two genomic regions) begins with the identification of similarity of a particular sequence or gene, but then requires additional support from flanking genomic regions or genes to identify correlated gene arrangements within a particular window size. The presence of flanking genomic regions with a high degree of synteny means that shared ancestry can be inferred with greater confidence for the gene pairs that reside in these regions. Users can inquire one gene from one species and investigate its syntenic homologs present in other species by presenting results of a certain window size (number of linking genes). However, these analyses are usually presented as linear aligned syntenic regions from limited number of species (Figure 1a). Other approaches for understanding and displaying relationships between homologous genes within and between different species include phylogenetic tree reconstructions and presence/absence scoring by phylogenetic profiling (Pellegrini et al. 1999, Kensche et al. 2008), but these approaches do not consider genomic context.

It is thus now possible to investigate syntenic relationships of all genes in a gene family context across a wide range of land plant species to address fundamental questions on the origin of novel gene function in relation to morphological changes and adaptations to niches. With continuous progress in sequencing technology and the sequencing of more plant species (such as gymnosperms and lower plants) the performance of our synteny network analysis will continue to improve.

By analyzing synteny networks, we uncover three cases of tandem gene arrangements of phylogenetically separated clades of MADS-box genes: *SEP1-SQUA*, *SEP3-SFT*, and *AGL6-TM3* in flowering plants over long evolutionary history (Figures 3 and 4). Moreover, together with MADS-box genes phylogeny, we were able to deduce the evolutionary history of the super- clade of MIKC^C^-type MADS-box genes. For the first time, we have also provided examples of angiosperm-wide conserved synteny networks of MIKC*-type genes (Supplemental Figures 1e and 1f), Type I MADS-box genes (Supplemental Figures 1g and 1h) and *StMADS11* (*SVP*) genes (Supplemental Figure 1c). Finally, we provide examples of lineage-specific networks derived from gene transpositions such as *AP3* genes (Figure 2d), *PI* genes (Figure 2e), and *FLC* genes (Supplemental Figure 1b) in Brassicaceae, and *AGL17* genes in monocot species (Supplemental Figure 1d).

In plants, the MADS-box genes are the major transcription factors that regulate the specification of floral organs, and they play a role in processes throughout the plant lifecycle. However, rarely have MADS-box genes been identified in regulatory clusters like animal Hox genes. This could be due more to the analysis techniques employed to date, namely phylogenetic analyses and pairwise synteny analyses, where ancient WGDs can dramatically complicate the analyses. The use of synteny networks to study evolutionary dynamics of plant gene families was previously done for the group VI ethylene responsive factors (van Veen et al., 2014) and more recently for the small ubiquitin-like modifiers gene family (Hammoudi et al., 2016). In contrast to these studies, our study represents a methodological roadmap of synteny network construction and analysis pipeline, which can then be applied to any gene families across many genomes. As the example, our results have provided insight into the MADS-box gene evolution at whole gene family level and generated new hypotheses on the birth and neofunctionalization of these genes during plant evolution.

## METHODS

### Plant Genomes Analyzed

In total, fifty-one plant genomes were included in our analysis (Supplemental Table 1 for detailed information), including thirty rosids, five asterids, *Beta vulgaris* (non-rosid non-asterid), eleven monocots, the early diverging angiosperm (*Amborella trichopoda*), and a single genome for gymnosperms (*Picea abies*), club moss (*Selaginella moellendorffii*), moss (*Physcomitrella patens*), and green alga (*Chlamydomonas reinhardtii*).

### Syntenic Block Calculation

MCScanX (Tang et al., 2008b; Wang et al., 2012) was used to compute genomic collinearity between all pairwise genome combinations using default parameters (minimum match size for a collinear block = 5 genes, max gaps allowed = 25 genes). The output files from all the intra- and inter- species comparisons were integrated into a single file named “Total_Synteny_Blocks”, including the headers “Block_Index”, “Locus_1”, “Locus_2”, and “Block_Score”, which served as the database file.

### Synteny Network for the MADS-box gene family

Candidate MADS-box genes were initially identified using HMMER3.0 with default settings (domain signature PF00319) (Finn et al., 2011) for each of the 51 genomes (Supplemental Table 2, sheet 1). Then this gene list containing all candidate MADS-box genes was queried against the “Total_Synteny_Blocks” file. Rows containing at least one MADS-box gene were retrieved into a new file termed “Syntenic_Blocks_MADS-box genes” (Supplemental Table 2, sheet). This file was then the final synteny network for the MADS-box genes, the network was imported and visualized in Cytoscape version 3.3.0 and Gephi 0.9.1 (Shannon et al., 2003).

Sequences were labeled based on the *A. thaliana* MADS-box genes plus three representative MADS-box genes that are not represented in *A. thaliana* (*TM8*-gene (GenBank Accession No. NP_001234105) from *Solanum lycopersicum*, *OsMADS32* gene (GenBank Accession No. XP_015642650) from rice, and *TM6* (GenBank Accession No. AAS46017) from *Petunia hybrida*) (Blanc and Wolfe, 2004; Lee et al., 2003; Daminato et al., 2014), using BLASTP (Altschul et al., 1990).

### Network Clustering

Clique percolation as implemented in CFinder (Derenyi et al., 2005; Palla et al., 2005; Fortunato, 2010) was used to locate all possible *k*-clique-communities for the MADS-box gene synteny network to identify communities (clusters of gene nodes). Increasing *k* values make the communities smaller and more disintegrated but also at the same time more connected. In this study, we used *k* = 6 for the networks represented in Figure 2e and the networks for phylogenetic profiling (Figure 2f), while in Figures 3a, 3b and S1, *k* = 3 was used for a more comprehensive analysis.

### Phylogenetic Profiling of Clustered Communities

Communities (synteny clusters) derived from a certain *k* value were extracted and the node (i.e. gene) composition of each community was then mapped to the phylogenetic tree with 51 species (Smith et al., 2011). Presence (red) or absence (white) of homologs in a particular cluster was depicted for the different species in the phylogenetic tree, thus creating a phylogenetic profile of a synteny cluster (Figure 2f). Each column in the illustration represents one community (one synteny cluster), which is labeled at top of the x-axis based on its MADS-box name/annotation. Through such clustering and phylogenetic profiling steps, representative communities for the Type II (MIKC^C^- and MIKC*- type) and Type I MADS box clades were found and then further analyzed.

### Phylogenetic Distance and Tree Construction

Amino acid sequences for the candidate MADS-box genes, both the genes represented in the synteny networks and the genes missing from the networks, were aligned using HmmerAlign (Kristensen et al., 2011). The alignment was then transferred into codon alignment using Pal2nal (Suyama et al., 2006). A phylogenetic tree and pairwise distance was computed using RAxML (Stamatakis, 2014) with the GTRCAT and GTRGAMMA model, respectively (bootstrap = 100). The depicted color key scale was generated based on the distribution of pairwise phylogenetic distances (Supplemental Figure 1a). The phylogenetic tree was annotated and depicted using iTOL v3 (Letunic and Bork, 2016).

## Supplemental Data

**Supplemental Table 1.** Genomes used in this analysis (including the tree used in Figure 3e with full species names and known duplication events)

**Supplemental Table 2.** Candidate MADS-box genes (sheet1) and synteny network for MADS-box genes (sheet2)

**Supplemental Table 3.** Communities at *k* = 6

**Supplemental Table 4.** Detail information for tandem arrangements

**Supplemental Figure 1.** Synteny network of other MADS-box gene clades

(a) Distribution of all pair wise phylogenetic distances of MADS-box genes. (b) Syntenic gene cluster for the Brassicaceae and Cleomaceae *FLC*-like genes. (c) *AGL17*-like syntenic gene cluster shared by monocot species. (d) Synteny network of the *StMADS11*- and *AGL17*-like genes. (e) Synteny network of the MIKC*-P clade genes. (f) Synteny network of the MIKC*-S clade genes. (g) Synteny network of the Type I-Mγ genes. (h) Synteny network of Type I-Mα genes.

## ACKNOWLEDGEMENTS

T.Z. was is supported by the China Scholarship Council (CSC), HvdB and MES by a Netherlands Scientific Organisation (NWO) Vernieuwingsimpuls Vidi grant (numbers 864.10.004 and 864.10.001, respectively). SdB received a NWO Experimental Plant Science graduate school “master talent” fellowship.

## AUTHOR CONTRIBUTIONS

MES and HvdB designed the research. T.Z. performed the analysis. R.H. and SdB analyzed data. T.Z. and MES wrote the article. All the authors critically read and commented on the article and approved of its final version for submission.

